# Suppression of HIV and cocaine-induced neurotoxicity and inflammation by cell penetrable itaconate esters

**DOI:** 10.1101/2023.09.25.559154

**Authors:** B. Celia Cui, Marina Aksenova, Aliaksandra Sikirzhytskaya, Diana Odhiambo, Elizaveta Korunova, Vitali Sikirzhytski, Hao Ji, Diego Altomare, Eugenia Broude, Norma Frizzell, Rosemarie Booze, Michael D. Wyatt, Michael Shtutman

**Affiliations:** Department of Drug Discovery & Biomedical Sciences, College of Pharmacy, University of South Carolina, Columbia, SC 29208, USA; Department of Pharmacology, Physiology & Neuroscience, School of Medicine, University of South Carolina, Columbia, SC 29208, USA; Department of Psychology, College of Arts and Sciences, University of South Carolina, Columbia, SC 29208, USA

**Author notes:** Authors contributed equally (B. Celia Cui, Marina Aksenova). Address correspondence to: M. Shtutman, 715 Sumter St, Columbia, SC, 29208, USA.

## Abstract

HIV-associated neurological disorder (HAND) is a serious complication of HIV infection, marked by neurotoxicity induced by viral proteins like Tat. Substance abuse exacerbates neurocognitive impairment in people living with HIV. There is an urgent need for effective therapeutic strategies to combat HAND comorbid with Cocaine Use Disorder (CUD). Our analysis of the HIV and cocaine-induced transcriptomes in primary cortical cultures revealed a significant overexpression of the macrophage-specific gene, aconitate decarboxylase 1 (Acod1), caused by the combined insults of HIV and cocaine. ACOD1 protein converts the tricarboxylic acid intermediate cis-aconitate into itaconate during the activation of inflammation. The itaconate produced facilitates cytokine production and subsequently activates anti-inflammatory transcription factors, shielding macrophages from infection-induced cell death. While the role of itaconate’ in limiting inflammation has been studied in peripheral macrophages, its immunometabolic function remains unexplored in HIV and cocaine-exposed microglia. We assessed in this model system the potential of 4-octyl-itaconate (4OI), a cell-penetrable esterified form of itaconate known for its potent anti-inflammatory properties and potential therapeutic applications. We administered 4OI to primary cortical cultures exposed to Tat and cocaine. 4OI treatment increased the number of microglial cells in both untreated and Tat±Cocaine-treated cultures and also reversed the morphological altercations induced by Tat and cocaine. In the presence of 4OI, microglial cells also appeared more ramified, resembling the quiescent microglia. Consistent with these results, 4OI treatment inhibited the secretion of the proinflammatory cytokines IL-1α, IL-1β, IL-6, and MIP1-α induced by Tat and cocaine. Transcriptome profiling further determined that Nrf2 target genes such as NAD(P)H quinone oxidoreductase 1 (Nqo1), Glutathione S-transferase Pi (Gstp1), and glutamate cysteine ligase catalytic (Gclc), were most significantly activated in Tat-4OI treated cultures, relative to Tat alone. Further, genes associated with cytoskeleton dynamics in inflammatory microglia were downregulated by 4OI treatment. Together, the results strongly suggest 4-octyl-itaconate holds promise as a potential candidate for therapeutic development aimed at addressing HAND coupled with CUD comorbidities.

Graphical Abstract:
Model of 4OI-mediated neuroprotection against Tat-Cocaine toxicity Tat and Tat-Cocaine treatment induce neuronal damage, which is mitigated by 4OI through microglia cells. This cartoon shows the reduction of harmful effects such as proinflammatory cytokine release, upregulation of P2R, PDE, and Acod1 by the presence of 4OI. This ester modified itaconate triggers anti-inflammatory responses and activates antioxidant pathways.

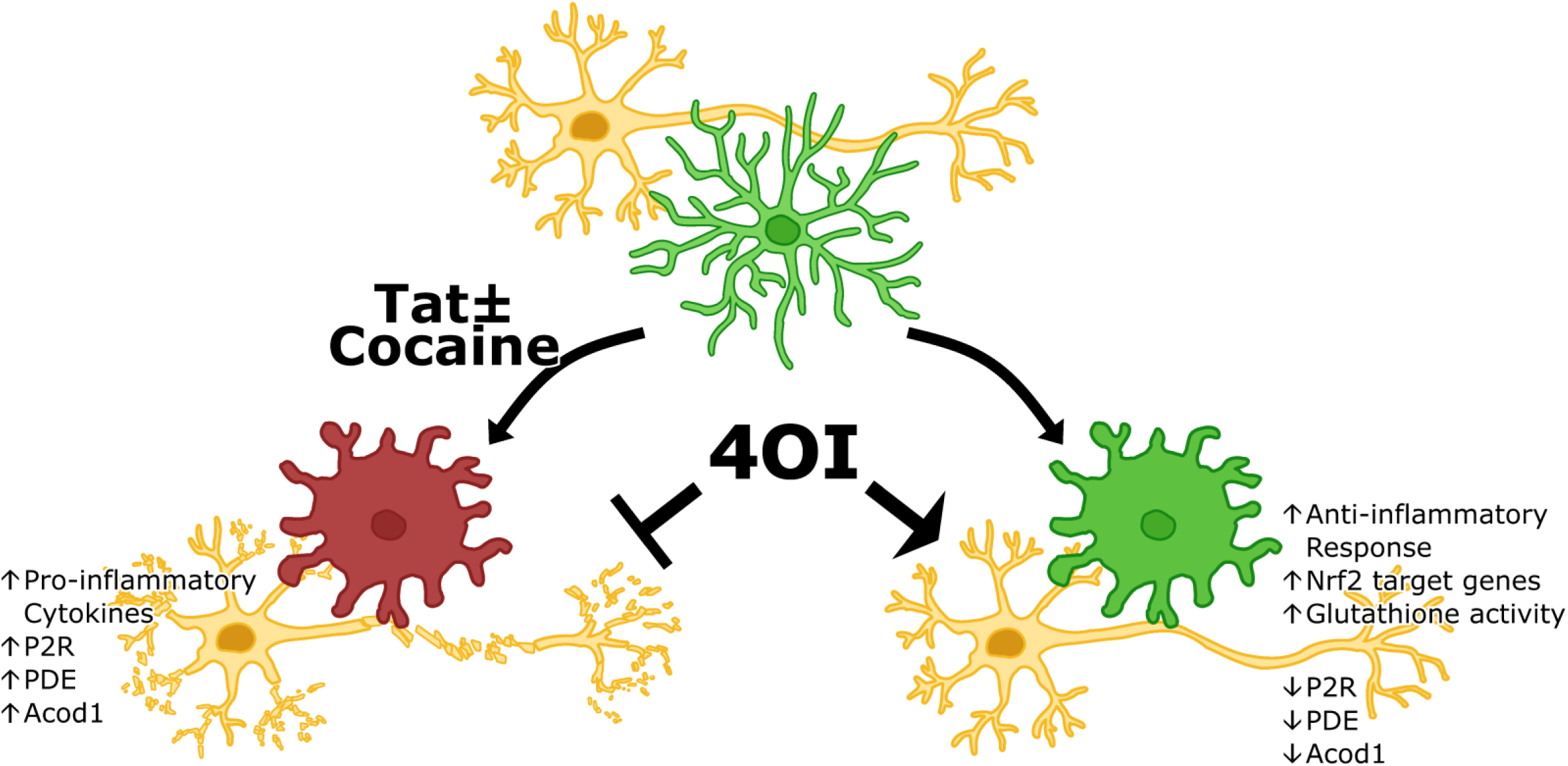

## Introduction

Human immunodeficiency virus type 1 (HIV-1) affects over 38 million individuals globally and over 1.2 million people in the United States (NIDA). The rise of antiretroviral therapy (ART) has effectively suppressed viral replication in people living with HIV (PLWH) and made the life expectancy of PLWH comparable to that of individuals without HIV infection. Nonetheless, PLWH continue to bear a disproportionately heavier burden of major comorbidities, experiencing these health challenges more than 16 years prior to the general population (Marcus, Leyden et al. 2020).

One of the most debilitating complications of HIV infection results in neurocognitive dysfunction referred to as HIV-associated neurological disorder (HAND). Despite the advent of ART, nearly 50% of PLWH continue to suffer from HAND. This condition encompasses a spectrum of neurological dysfunctions, spanning asymptomatic neurocognitive impairment and mild neurocognitive disorder, to HIV-associated dementia, which results in severe cognitive deficits and memory loss (Saylor, Dickens et al. 2016). Given the considerable burden on affected individuals’ quality of life, it is crucial to alleviate the neurotoxicity of HIV infection in the cells of the central nervous system (CNS). Microglial cells, despite being only 15% of total brain cells, play a vital role in HIV infection within the CNS. They serve as the primary site for viral infection and replication, as well as the main reservoir and source of viral latency in the brain. Moreover, microglia contribute significantly to HIV reactivation and the development of a chronic cerebral inflammatory state (Wallet, De Rovere et al. 2019, Sreeram, Ye et al. 2022).

Reportedly, all HIV proteins, including transactivator of transcription (Tat), gp120, gag, env, have been shown to exert neurotoxic effects on the progression of CNS pathology (Nath 2002, Yuan, Huang et al. 2015). Among these proteins, Tat is considered one of the most toxic, and functions as an early regulatory protein essential for the replication capacity of the virus. Tat mediates HIV gene expression and controls the complex pattern of protein expression necessary for functional viral transcription (Karn and Stoltzfus 2012). Additionally, Tat is highly implicated in intracellular pathways that lead to significant proinflammatory cytokine overproduction in macrophages (Ben Haij, Planes et al. 2015, Zayyad and Spudich 2015).

Compounding the HIV-associated neuropathology, there is a strong correlation between substance use disorders and the exacerbation of HAND (Beyrer, Wirtz et al. 2010, Gannon, Khan et al. 2011). Notably, cocaine, one of the most abused drugs in the US (NIDA 2018), is frequently used among PLWH. The concomitant use of HIV and cocaine has been shown to significantly contribute to HIV pathogenesis. Adding to this, HIV-infected patients with cocaine use disorder (CUD) often experience worsened HAND (Avants, Margolin et al. 1997, Atluri 2016). Previous studies have revealed that cocaine can activate immune responses and induce the expression of proinflammatory cytokines from BV2 microglia cells (Liao, Guo et al. 2016). While the combined consequences of Tat and cocaine are multifaceted, our specific focus centers on understanding how the interactions between Tat and cocaine contribute to the induction of neuroinflammation through microglia (Zhao, Zhang et al. 2022).

Current ART drugs struggle to adequately penetrate blood brain barrier (BBB), are prone to effective removal from brain parenchyma, and as a result, fail to combat HIV reservoir formation and persistence in the brain (Osborne, Peyravian et al. 2020). Regrettably, there is currently no approved therapy available for the treatment of HAND, especially for the neurological consequences resulting from concurrent HIV infection and drug abuse. However, targeting aberrant microglial activity has emerged as a potential therapeutic approach for treating both neurogenerative diseases and drug addiction (Biber, Bhattacharya et al. 2019, Catale, Bussone et al. 2019, Liu, Wang et al. 2019). As a result, microglia infected with the virus and the resulting inflammatory response triggered by the toxic HIV proteins are increasingly recognized as a disease-associated microglial state that requires further investigation. Previously, we have uncovered that the inhibition of microglial activation via a small molecule inhibitor can protect neurons from the combined Tat and cocaine toxicity, suggesting that microglia can be a valuable target in HAND-CUD treatment (Aksenova, Sybrandt et al. 2019).

We observed an increased expression of aconitate decarboxylase 1 (Acod1) based on RNA sequencing analysis of Tat-and cocaine-treated primary cortical cultures. Acod1, also known as immune responsive gene 1 (IRG1), catalyzes the synthesis of itaconate by decarboxylating the tricarboxylic acid cycle intermediate cis-aconitate, which shunts this carbon skeleton out of the TCA cycle (Michelucci, Cordes et al. 2013). Itaconate can be added to protein thiols through a process called dicarboxypropylation and alter their activity or function. Notably, itaconate is emerging as an essential regulator of immune metabolism, displaying high levels in activated macrophages and influencing in their metabolic functions and inflammatory responses (O’Neill and Artyomov 2019). Considering that HIV-induced metabolic dysfunction is exacerbated by cocaine (Samikkannu, Atluri et al. 2016), itaconate becomes even more intriguing in the context of this virus.

Recent studies have investigated the immunomodulatory properties of 4-octyl-itaconate (4OI), a membrane and BBB penetrable ester-modified itaconate. In particular, 4OI has demonstrated potent anti-inflammatory effects and has been shown to significantly reduce inflammatory responses to a variety of pathogenic viruses such as SARS-CoV-2, Herpes Simplex Virus-1 and -2, Zika and Vaccinia (Olagnier, Farahani et al. 2020, Sohail, Iqbal et al. 2022). The anti-inflammatory effect of itaconate is mediated in part by its ability to regulate immune function and inflammation by delaying pyroptotic cell death (Bambouskova, Potuckova et al. 2021), attenuating the excessive production of reactive oxygen species (Sohail, Iqbal et al. 2022), promoting anti-inflammatory transcription factor Nrf2 through Nrf2-Keap1 signaling pathway (Mills, Ryan et al. 2018, Ni, Xiao et al. 2022), and inhibiting pro-inflammatory cytokine production (Lampropoulou, Sergushichev et al. 2016).

However, the role of the Acod1/itaconate axis in HIV-1 and cocaine-induced neurotoxicity is unknown. To investigate, we tested the effects of the cell-permeable itaconate derivative 4OI in Tat and cocaine-treated primary rat fetal cortical cultures. Our study reveals, for the first time, that 4OI can provide neuroprotection against Tat and cocaine neurotoxicity in vitro. Additionally, we observed 4OI-induced changes in microglia and conducted transcriptomic analyses on these cell cultures to obtain a better understanding of the underlying mechanisms.

## Materials and Methods

### Neuronal Cell Cultures

Primary rat fetal mixed neuronal cultures were prepared from 18-day-old Sprague-Dawley rat fetuses (Envigo Laboratories, Indianapolis, IN) as described previously (Aksenova, Aksenov et al. 2009, Aksenova, Sybrandt et al. 2019) in accordance with the University of South Carolina Institutional Animal Care and Use Committee. Briefly, tissue was dissected, incubated in a solution of 0.05% Trypsin/EDTA, washed with Hank’s balanced salt solution (HBSS, Thermo Fisher Scientific), dissociated by trituration and distributed into poly-L-lysine coated 12-well plates (Costar, Cambridge, MA) with or without inserted glass coverslips containing DMEM/F12 medium with 10% fetal bovine serum. After 24 hours initial plating medium was replaced with serum-free Neurobasal medium supplemented with 2% B-27, 2 mM GlutaMAX and 0.5% D-glucose. All reagents were from Thermo Fisher Scientific. Half of the medium was replaced with freshly prepared medium once a week. Cultured cells were used for experiments at the age of 3 weeks in vitro (DIV 21).

### Experimental treatment

Primary rat cortical cultures were treated with recombinant Tat 1-86 (Diatheva, Italy) and with cocaine-HCl (Sigma Chemicals) as described in each Figure Legend. The concentrations of Tat and cocaine were selected in accordance with our previous experimental data (Aksenova, Sybrandt et al. 2019), and stock solutions were prepared as previously described. 4-octyl-itaconate was obtained from Cayman Chemicals (Catalog No. 25374, Ann Arbor, MI). A stock solution of itaconate was prepared as a 50 mM stock in DMSO and was diluted to final concentrations from 30 μM up to 250 μM.

The concentration of Tat used in our study was chosen to reflect those found in the sera of HIV-positive patients (Xiao, Neuveut et al. 2000) and in cerebrospinal fluid (Westendorp, Frank et al. 1995). Similarly, the concentrations of cocaine used in the study were chosen based on animal studies that involved drug self-administration, which were suggested to parallel those found in recreational human users (Zimmer, Dobrin et al. 2011) and postmortem brain tissues in fatal cases of cocaine abuse (Spiehler and Reed 1985)

### Apoptotic/dead cells detection

Dead and apoptotic cells were detected using the CellEvent Caspase-3/7 Kit (#C10423, Thermo Fisher Scientific) according to the manufacturer’s recommendations. Briefly, after experimental treatment, Caspase3/7 Green Detection Reagent was added directly to cells, and the plate was incubated for 30 min at 37 °C. During the final 5 min of incubation, SYTOX AADvanced dead cell solution was added. Cells were rinsed with PBS, and images of cells were taken immediately. Alternatively, cells were fixed with 4% paraformaldehyde, imaged, and used for further experiments.

### Immunocytochemistry

For ICC analysis cells were plated on glass coverslips and placed inside 12-well plates. Following experimental treatment, primary cortical cultures were fixed with 4% paraformaldehyde (EM Sciences, Hatfield, PA) and permeabilized with 0.1% Triton X-100. Fixed cultures were blocked with 10% fetal bovine serum for 2 h and then co-labeled overnight with the following primary antibodies: chicken polyclonal anti-MAP2 antibodies (1:2,000) (# ab92434, Abcam, Cambridge MA) or rabbit monoclonal anti-Iba1 (1:300) (# ab178847 Abcam, Cambridge MA). Secondary antibodies, goat anti-chicken IgG conjugated with AlexaFluor 594, and goat anti-rabbit IgG conjugated with AlexaFluor 488 (1:500 both; Invitrogen Life Technologies, Grand Island NY), were used for visualization. To identify cell nuclei, DAPI was added with the final PBS wash, and coverslips were mounted on glass slides using ProLong Glass Antifade Mount (Invitrogen Life Technologies, Eugene, OR).

### Image processing and analysis

Images were captured on a Carl Zeiss LSM 700 laser scanning confocal microscope (Carl Zeiss Meditec, Dublin, CA) equipped with 20x (Plan APO 0.8 air) objective. Images were captured using 0.7 scanning zoom with 312-nm X-Y pixel size. Fluorescence and differential interference contrast (DIC) imaging was done using single-frame mode.

ImageJ software (National Institutes of Health, USA) was used for manual analysis of microscopy images acquired using a Zeiss 700 confocal microscope. Several fields of vision were taken from at least three different wells. The total number of cells and percentage of Iba1 positive cells were estimated using segmentation of DNA channel (DAPI) followed by “Analyze Particles” ImageJ command. Size of microglial cells and length of microglia processes were estimated individually using “Freehand selections” and “Freehand lines” ImageJ tool. Data were aggregated, analyzed and visualized using R ggplot2 tools. Background correction of widefield images was performed by background (Gaussian blur) division procedure (32-bit mode) followed by image histogram adjustment for 16-bit dynamic range.

### Cytokine/chemokine array

Medium was collected from each culture well at the end of each experiment, frozen, and sent to Eve Technologies Corporation (Calgary, Canada) for LUMINEX-based analysis of cytokines by Featured-Rat Cytokine Array/Chemokine Array 27-plex (RD27). The 27-plex array analyzed Eotaxin, EGF, Fractalkine, IFN-γ,IL-1α, IL-1β, IL-2, IL-4, IL-5, IL-6, IL-10, IL-12 (p70), IL-13, IL-17A, IL-18, IP-10, GRO/KC, TNF-α, G-CSF, GM-CSF, MCP-1, Leptin, LIX, MIP-1α, MIP-2, RANTES, VEGF.

### RNA sequencing

RNA and library preparation, post-processing of the raw data and data analysis were performed by the USC CTT COBRE Functional Genomics Core. RNAs were extracted using Zymo Quick-RNA MicroPrep Kits as per manufacturer recommendations (Zymo Research, Irvine, CA, USA). RNA quality was evaluated on RNA-1000 chips using a Bioanalyzer (Agilent, Santa Clara, CA, USA). RNA libraries were prepared using an established protocol with NEBNExt Ultra II Directional Library Prep Kit (NEB, Lynn, MA). Each library was made with one of the TruSeq barcode index sequences and the Illumina sequencing done by Medgenome (Foster City, CA) with Illumina NovaSeq PE100 (100bp, pair-ended). Sequences were aligned to the Rat genome Rnor_6.0 (GCA_000001895.4, ensemble release-99) using STAR v 2.7.2b (Dobin, Davis et al. 2013). Samtools (v1.2) was used to convert aligned sam files to bam files and reads were counted using the featureCounts function of the Subreads package (Liao, Smyth et al. 2014) and the Rattus_norvegicus.Rnor_6.0.93.gtf annotation file. Only reads that were mapped uniquely to the genome were used for gene expression analysis. Differential expression analysis was performed in R using the edgeR package (Robinson, McCarthy et al. 2010). Raw counts were normalized using the Trimmed Mean of M-values (TMM) method and the normalized read counts were then fitted to a generalized linear model using the function glmFit (McCarthy, Chen et al. 2012). Genewise tests for significant differential expression were performed using the function glmLRT. The P-value was then corrected for multiple testing using Benjamini-Hochburg’s FDR.

### Data analysis and statistics

Graphs were generated using Inkscape version 1.3. Data visualization and analysis were performed using R v4.3.1 with ggplot2 v3.4.3 in RStudio v2023.06.1+524 using custom code. The Mann-Whitney-Wilcoxon Test followed by Bonferroni correction were utilized for statistical comparisons of relevant experimental groups, unless specified otherwise. All data are represented as mean ± SD unless otherwise stated, with p<0.05 regarded as statistically significant.

## Results

### Transcriptomic profiles of Tat and cocaine treated primary cortical cultures reveals differential expression of Acod1

To investigate the impact of Tat and cocaine on the transcriptome of primary cortical cultures, we previously performed RNA sequencing analysis on cultures treated with Tat alone, cocaine alone, or a combination of both (Aksenova, Sybrandt et al. 2019). Analysis of the transcriptome upregulated by Tat or Tat-Cocaine treatment revealed a significant increase in the expression of Acod1 (Figure 1A). To corroborate this finding, we performed qPCR analysis measuring the expression of Acod1 normalized to the reference control ACTB expression within the same experimental cultures. The qPCR results confirmed a significant increase in Acod1 expression relative to the control for Tat only or Tat and cocaine treatment groups (Figure 1B).

**Figure 1:**
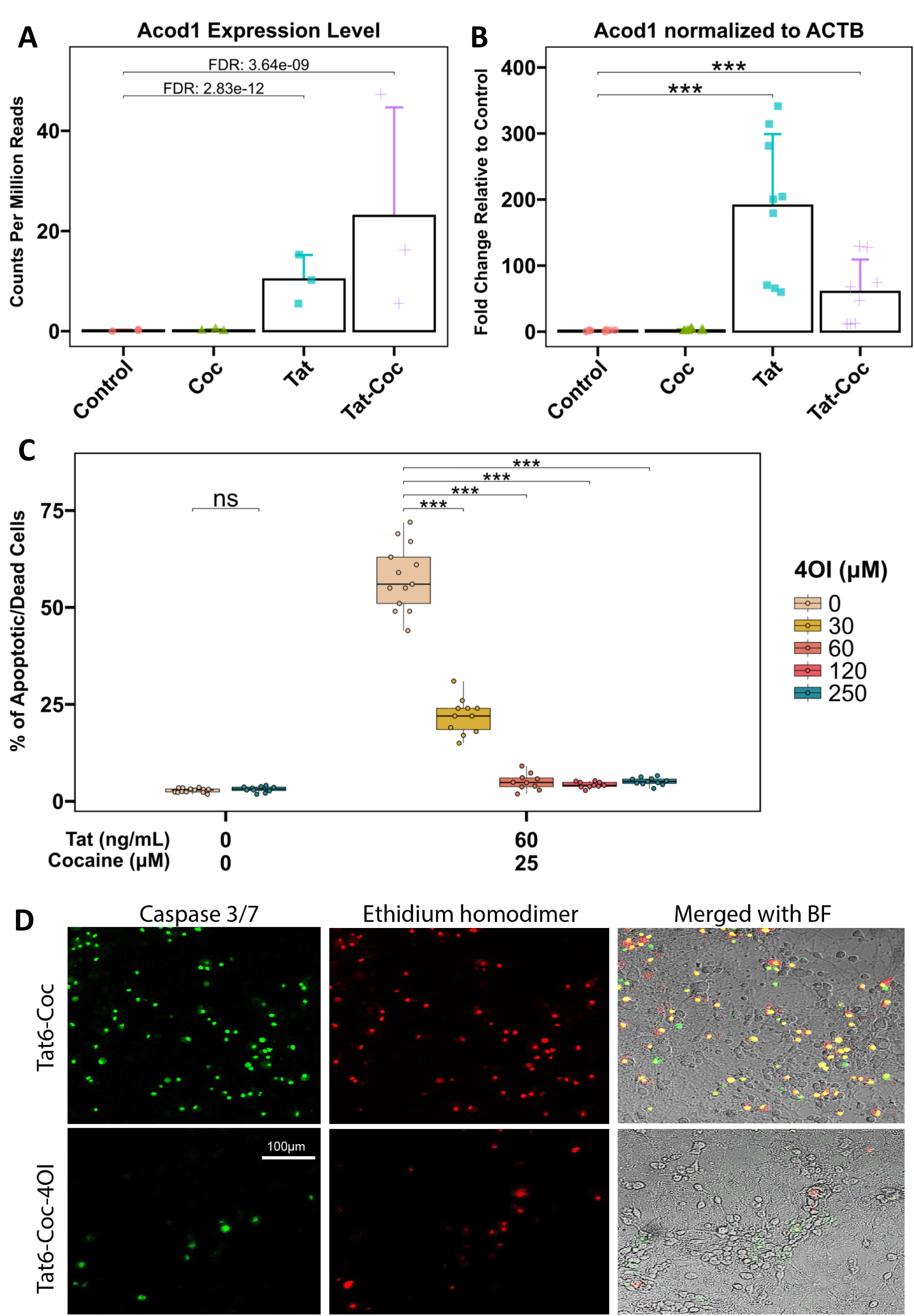
Acod1 expression in Tat and/or cocaine-treated primary neuronal cultures, and neuroprotective effects of 4OI. **(A)** Acod1 expression was assessed in primary cortical cultures treated with cocaine (25 μM) alone for 24 h, Tat (6 ng/mL) alone for 48 h, or Tat for 48 h followed by the addition of cocaine for 24 h. Acod1 levels were quantified using RNA-seq and presented as CPM read counts. **(B)** qPCR analysis of the same experimental samples measured Acod1 expression fold change, normalized to ACTB. Statistical analysis was performed using the Mann-Whitney/Wilcoxon Rank-Sum test, followed by Bonferroni’s post hoc test. **(C)** Immunofluorescent staining of cultures treated with Tat (60 ng/mL) with or without 4OI (250 μM) for 48 h followed by cocaine (25 μM) for 24 h, fixed with 4% PFA. Dead/apoptotic cells were detected using the CellEvent Caspase 3/7 Assay Kit (Green: Caspase 3/7; Red: Ethidium Bromide). Scale bar: 100 μm. **(D)** Boxplots illustrate the dose-dependent effect of 4OI (30, 60, 120, or 250 μM) on Tat (60 ng/mL) and cocaine (25 μM) treated cultures. Non-treated control cultures and cultures treated with the highest concentration of 4OI (250 μM) are shown on the left. Each data point represents an image containing 300-450 cells. Statistical significance was determined using the Mann-Whitney-Wilcoxon test, followed by Bonferroni post hoc test (ns: not significant; *: p<0.05; **: p<0.01; ***: p<0.001).

Although the role of Acod1 in the context of HAND has not been examined, there is a rapidly emerging recognition of the importance of its product, the metabolite itaconate, in cellular inflammatory pathways and in the progression of various viral infections (Olagnier, Farahani et al. 2020, Sohail, Iqbal et al. 2022). In particular, the introduction of exogenous itaconate was shown to exhibit antiviral effects and to curtail inflammatory responses (Daniels, Kofman et al. 2019). Thus, we chose to evaluate the effects of itaconate in this HAND-CUD model.

### Neuroprotective effect of 4-octyl-itaconate against Tat and cocaine combined neurotoxicity in primary cortical cultures

After identifying Acod1 as a Tat and Tat-Cocaine upregulated gene, (Figure 1A-B), we evaluated the effects of itaconate, the product of cis-aconitate decarboxylation by Acod1. Because itaconate in its cytosolic form is not cell-permeable, we opted to use its membrane-penetrable ester-modified derivative, 4-octyl itaconate (4OI), to investigate the effects of metabolic modulation on microglia function in our HAND and CUD primary culture model that uses the Tat protein and cocaine as the neurotoxic insults. Consistent with our previous study, treatment of the cortical cultures with Tat (1, 6, or 60 ng/mL) or cocaine (10 or 25 μM) individually at the doses tested did not result in significant cell death, while the combination of Tat and cocaine resulted in concentration-dependent neuronal death (Supplemental Figure 1A). The combined neurotoxic effect was not observed when cortical cultures were treated with heat-inactivated Tat combined with cocaine.

The administration of 4OI successfully reversed the detrimental effects of the combined Tat and cocaine toxicity (Figure 1C-D, Figure 2), with the decrease in cell death due to 4OI exhibiting a dose-dependent response (Figure 1C). Based on these results, we identified 4OI doses of 60 μM and 250 μM as the lowest and highest concentrations that significantly rescued neuronal cell death under these conditions. These findings strongly suggested that treatment with 4OI can effectively rescue neurons from the death caused by the presence of Tat and cocaine.

**Figure 2:**
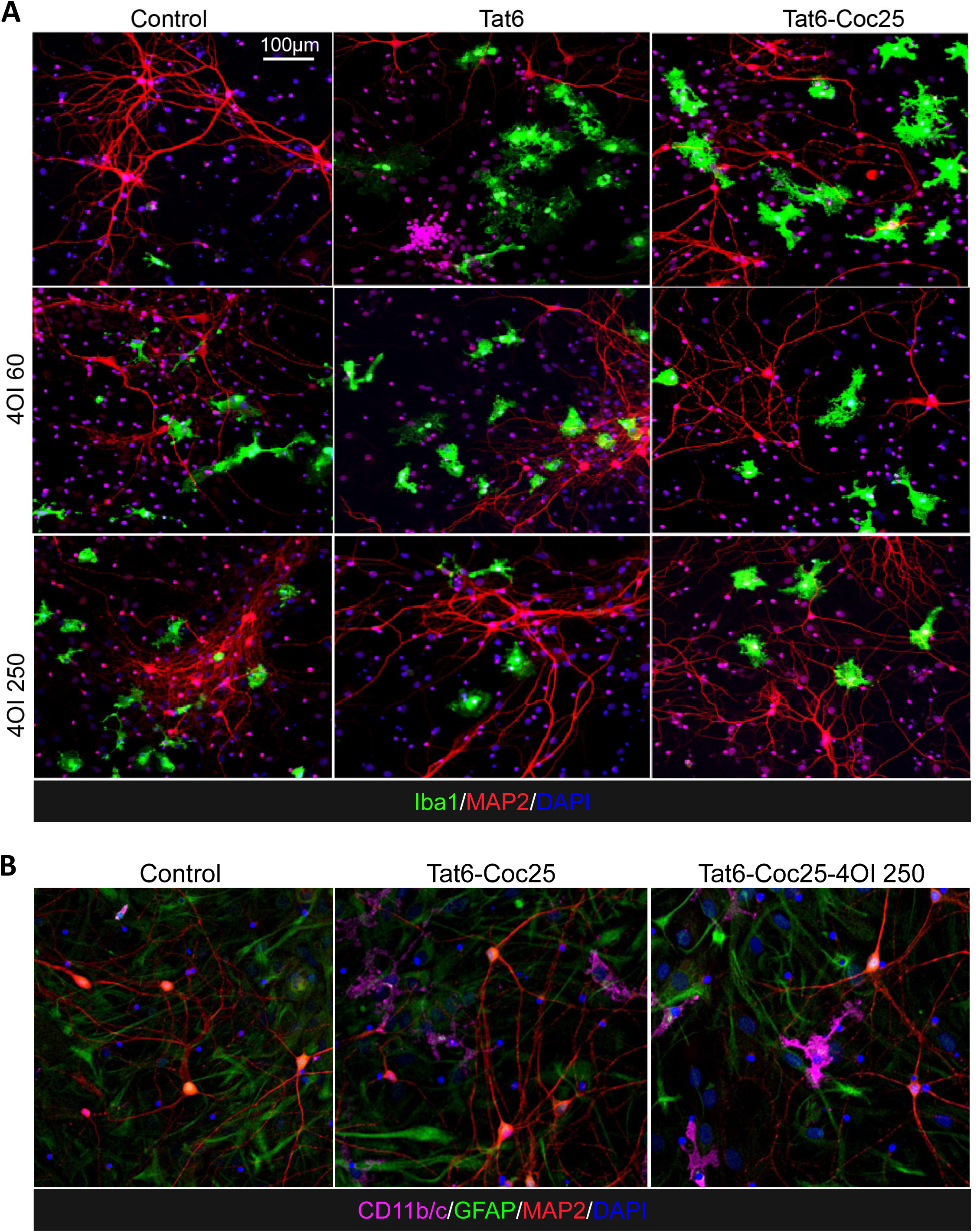
Effects of 4OI treatment on microglia morphology in Tat and/or cocaine-treated primary neuronal cultures. Immunofluorescent staining of primary cortical cultures: **(A)** Cells were treated with Tat alone (6 ng/ml) or Tat combined with 4OI (60 or 250 μM) for 48 h, followed by cocaine (25 μM) for additional 24 h. Cells were fixed with 4% PFA. Staining: Iba1 (green, microglia marker), MAP2 (red, neuronal marker), DAPI (blue, nuclei). Scale bar: 100 μm. **(B)** Cells were subjected to the same treatment. Staining: CD11b/c (magenta, activated microglia marker), GFAP (green, astrocyte marker), MAP2 (red, neuronal marker), DAPI (blue, nuclei).

### 4OI modulates the morphological changes of microglia brought by Tat and cocaine treatment

The neuroinflammation induced by Tat or Tat-Cocaine treatment appeared to manifest in morphological changes observed in microglia. Exposure to Tat alone or Tat-Cocaine resulted in an increased percentage of microglial cells (Supplemental Figure 1B), accompanied by a greater occurrence of microglia exhibiting amoeboid-shape characteristics (Supplemental Figure 1C-D). This ameboid shape is indicated by a reduction in process length and an increase in the cell body size of microglia. This effect was Tat and Tat-Cocaine concentration-dependent, except for the highest doses used in combination, which resulted in a decrease in the number of microglia cells and their body size commensurate with the increase in cell death at this highest combination dose (Supplemental Figure 1).

Treating naïve cultures with 4OI alone also resulted in an increase in microglial cells displaying more ameboid characteristics (Figure 3B-C). In particular, when cultures were treated with 250 μM 4OI (represented in dark blue), the median microglial processes shortened by about 5 μm, showing a 20% decrease, while the median size of microglial cell bodies expanded by about 250 μm², showing a 100% increase (Figure 3B-C). Interestingly, microglia morphological profiles in cultures treated solely with 4OI closely resembled the activated microglia states observed in Tat-Cocaine treated cultures. This similarity was evident in the 3D spread of data representing total quantified microglia morphology representing percentages of cells by size and length. Specifically, the expanded spherical shape of the 3D plot showed a greater resemblance between the Tat-Cocaine and 4OI samples, in contrast to the control spread, which took a flat disk shape (Figure 3D-E).

**Figure 3:**
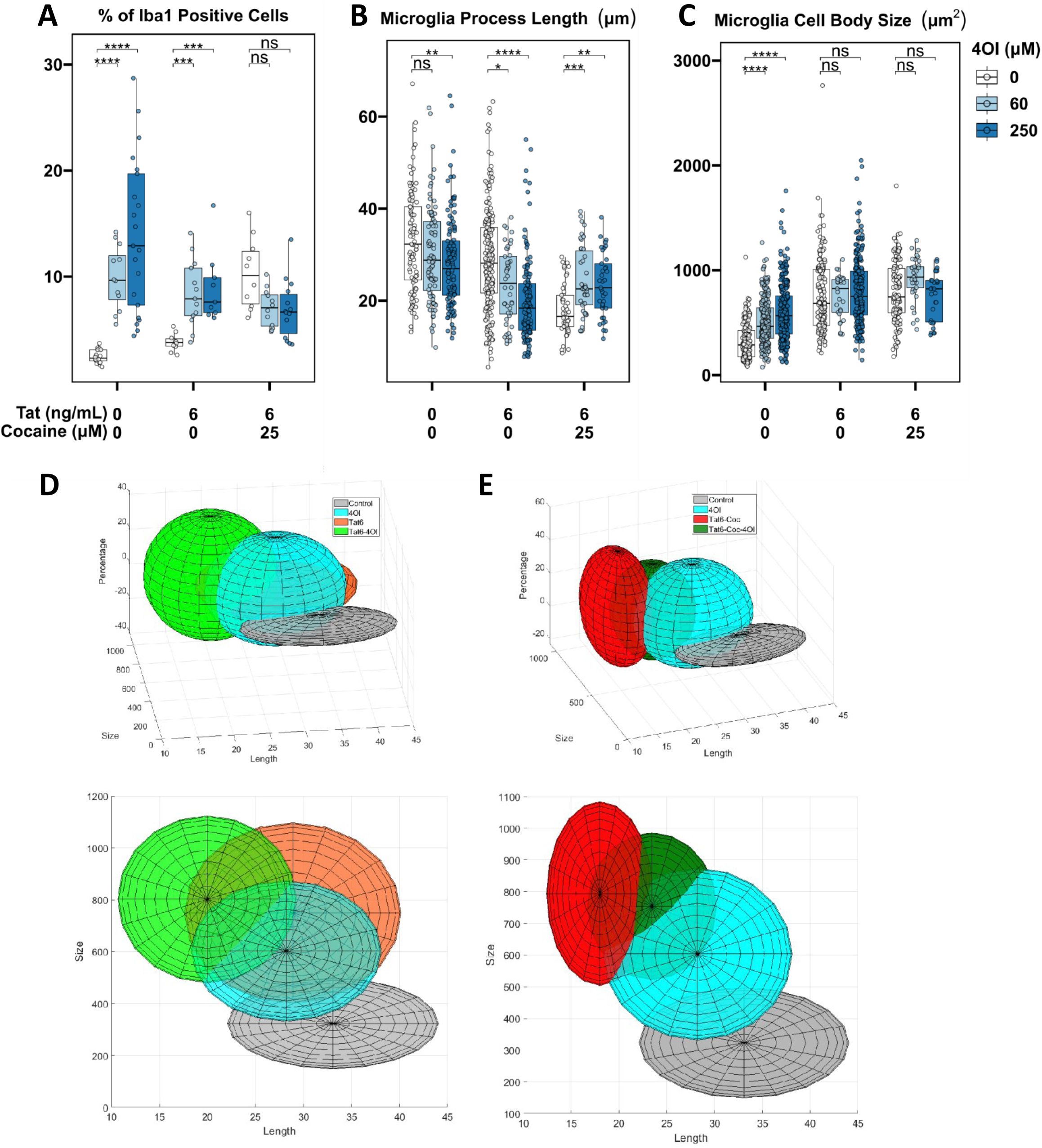
Effects of Tat and cocaine on microglia cell morphology in primary neuronal cultures. **(A)** Proportion of Iba1 positive microglial cells in cultures treated with Tat (6 ng/mL) and/or 4OI (60 or 250 µM) for 48 h, with or without an additional 24 h of cocaine (25 µM). Tat-Coc vs Tat-Coc-4OI (60 µM) yielded not significant p-adj value = 0.126. Tat-Coc vs Tat-Coc-4OI (250 µM) yielded not significant p-adj value = 0.114. **(B)** Length of microglia processes and (**C**) microglia cell body size in the same cultures. The intensity of the blue color indicates increasing 4OI concentration (60, 250 µM). The box covers 50% of the data in each condition, and the line inside indicates the median value. The Mann-Whitney-Wilcoxon test was conducted to calculate the statistical significance, followed by Bonferroni post hoc test (ns, not significant; *, p<0.05; **, p<0.01; ***, P<0.001). **(D)** 3D projection of microglia morphology affected by 4OI and/or Tat treatment. **(E)** 3D projection of microglia morphology affected by 4OI and/or Tat-Cocaine treatment (x: Body size; y: Percentage; z: Process length).

The effects of 4OI on microglia morphology were not unidirectional; instead, they were context dependent. The introduction of 4OI in Tat and cocaine treated cultures yielded opposing trends. For example, low Tat (6 ng/mL) and cocaine caused an increase in the percentage of Iba1-positive cells, along with a decrease in both microglia process length and cell body size, which were countered by 4OI treatment (Figure 3A-C). Similarly, the effect of high Tat (60 ng/mL) and cocaine, which led to reduced Iba1-positive microglial cell percentage, microglia process length, and cell body size, were countered by 4OI administration (Supplemental Figure 2A-C).

The 3D data spread illustrate that microglia treated with Tat-Cocaine-4OI were trending to the morphological state observed in 4OI-only treatments (Figure 3D-E). Therefore, while 4OI mitigated Tat-Cocaine toxicity, 4OI alone also drove microglia towards an amoeboid morphological state.

### RNA transcriptome of cell cultures treated with Tat, cocaine, and 4OI

To gain insights into the transcriptomic changes induced by Tat (6 ng/mL) or Tat and cocaine (25 μM) in primary cortical cultures with or without 4OI (250 μM) treatment, we conducted RNA-seq analysis (Supplemental Table 1). The bulk RNA sequencing findings were consistent with our earlier investigation (Aksenova, Sybrandt et al. 2019), revealing a substantial impact of Tat treatment on the transcriptome (Figure 4A). In contrast, the 24 h exposure to cocaine alone had a comparatively minimal effect on gene expression (Figure 4C). In accordance with our previous results, the combined treatment Tat and cocaine yielded outcomes that closely mirrored those observed with Tat alone (Figure 4B).

**Figure 4:**
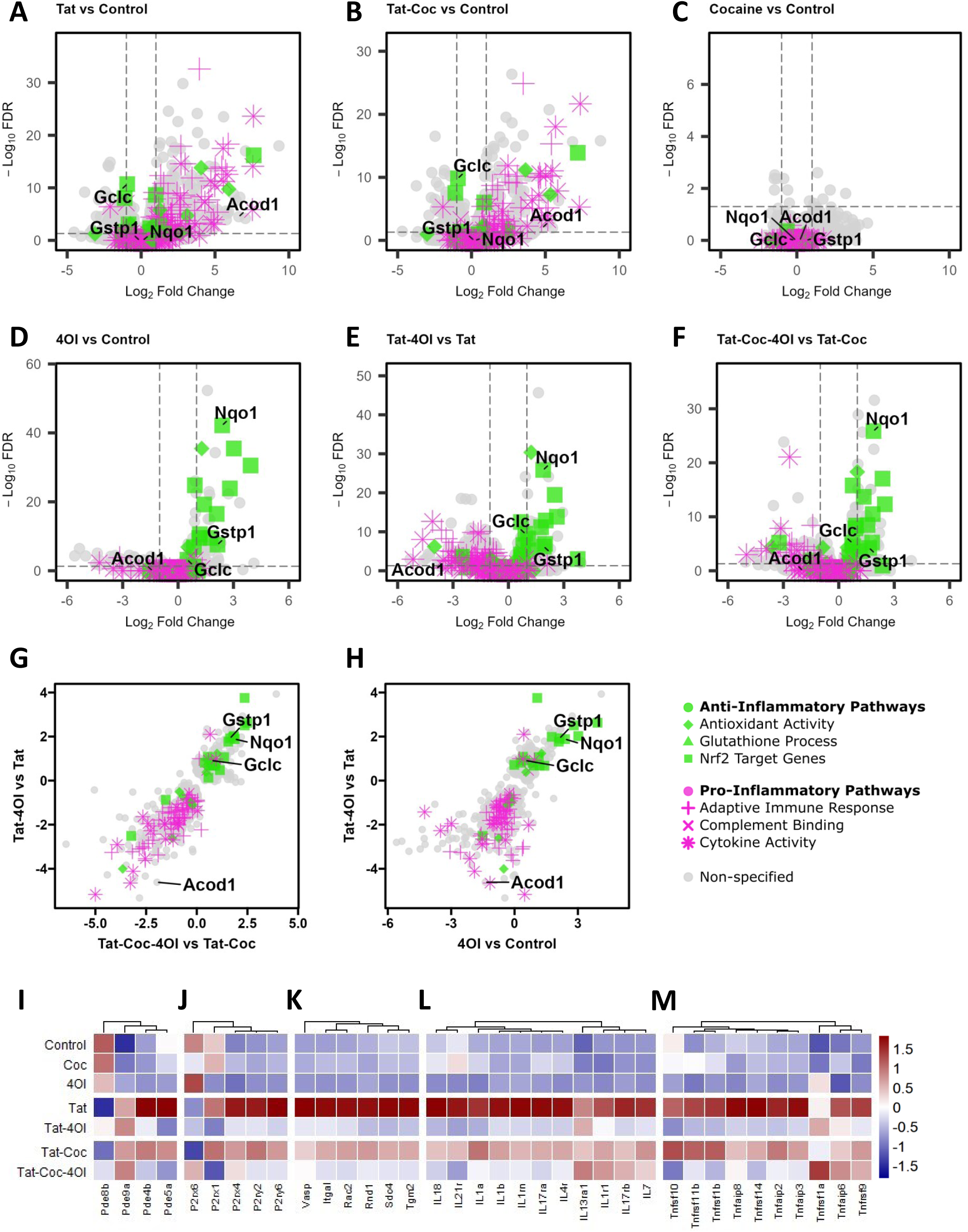
Genome-wide effects of 4OI on primary neuronal cultures treated with Tat and/or cocaine. RNA-seq transcriptomics of primary cortical cultures treated with Tat (6 ng/mL) and/or cocaine (25 μM) with/without 4OI (250 μM). Volcano plot comparing differential gene expression levels of (**A**) Tat treated samples vs Control, (**B**) Tat-Cocaine treated samples vs Control, (**C**) and Cocaine treated samples vs Control. Volcano plot comparing the differential gene expression between (**D**) 4OI vs Control (**E**) Tat-4OI vs Tat, and (**F**) Tat-Cocaine-4OI vs Tat-Cocaine conditions. Scatter plot illustrating the effect of 4OI on transcriptomic fold changes of select differentially expressed genes (FDR<0.05), and how they correlate between Tat-4OI vs Tat against (**G**) 4OI vs Control and (**H**) Tat-Cocaine-4OI vs Tat-Cocaine conditions. The colors correspond with the listed gene set pathway, with green representing select anti-inflammatory and magenta representing select pro-inflammatory pathways. Heatmap showing gene expression levels for (**I**) phosphodiesterase-associated genes, (**J**) purinergic receptor associated genes, (**K**) phagocytic cytoskeleton-associated genes, (**L**) interleukin superfamily genes, and (**M**) tumor necrosis factor family genes.

The pathway enrichment analysis of genes regulated by Tat alone or combined Tat-Cocaine exposure corroborated findings from other studies, demonstrating notable enrichment across diverse pathways. These included the adaptive immune response, complement binding, cytokine activity, and pathways regulated by NF-κB (Figure 4A-B). Treatment with 4OI alone or in conjunction with Tat or Tat-Cocaine dramatically induced the expression of Nrf2 target genes (Figure 4D-F, Table1-2). Remarkably, Nrf2 targeted pathways such as glutathione reduction antioxidant pathway involving NAD(P)H quinone oxidoreductase 1 (Nqo1), Glutathione S-transferase Pi (Gstp1), and glutamate cysteine ligase catalytic subunit (Gclc), displayed noteworthy activation exclusively in primary cortical cultures treated with 4OI (Figure 4A-F). 4OI administration led to an upregulation of anti-inflammatory pathways, including Nrf2 target genes and antioxidant activity, alongside a downregulation of pro-inflammatory pathways such as cytokine activity and adaptive immune response (Figure 4G-H). This enrichment of Nrf2 target gene pathways and antioxidant activities suggests a potential neuroprotective mechanism of 4OI in mitigating adverse neurotoxic effects.

Importantly, treatment with 4OI demonstrated a reduction in Tat-activated expression of well-known drivers of microglia inflammation, such as phosphodiesterase 4 (Pde4b) (Pearse and Hughes 2016) (Figure 4I), along with P2 purinergic receptors P2rx4, P2ry2, and P2ry6 (Anwar, Pons et al. 2020, Sophocleous, Ooi et al. 2022, Hide, Shiraki et al. 2023) (Figure 4J). These purinergic receptors are recognized regulators of active phagocytosis and removal of stressed neurons (Puigdellivol, Milde et al. 2021). Moreover, the actin-binding proteins that coordinate phagocytosis, notably Vasodilator Stimulated Phosphoprotein (VASP) (Montaño-Rendón, Walpole et al. 2022), showed upregulation due to Tat-Cocaine treatment and downregulation in the presence of 4OI treatment (Figure 4K). Similar trends were observed in the expression of proinflammatory cytokines, such as the interleukin and tumor necrosis factor families. Upregulation induced by Tat or Tat-Cocaine exposure was counteracted by 4OI administration, leading to downregulation (Figure 4L-M, Supplemental Figure 4).

Tat treatment led to a significant induction of Acod1 expression in comparison to the untreated control (Figure 1A-B). This upregulated Acod1 expression induced by Tat or Tat-Cocaine was markedly diminished upon 4OI administration (Figure 4E-F). Supported by additional qPCR analysis, the inclusion of 4OI exhibited a near three-fold reduction in Acod1 expression regardless of the presence of Tat or Tat-Cocaine, when compared to cultures treated with Tat alone (Supplemental Figure 3A-B). Collectively, transcriptomic analysis of the experimental samples unveiled distinct gene expression patterns, providing insights into the specific influence of 4OI in modulating neuroinflammatory pathways

### 4OI inhibition of proinflammatory cytokine secretion from Tat and cocaine treatment

We sought to uncover the functional consequences of 4OI administration on microglial cells in our primary mixed cultures, focusing on cytokine secretion as a key biomarker of inflammatory response. The release of proinflammatory cytokines serves as a widely recognized indicator of the detrimental effects associated with microglia activation (Hanisch 2002). To measure the levels of cytokines, we collected media from mixed cortical cultures treated with Tat followed by cocaine, with or without 4OI. We then determined the levels of 27 cytokines. The neuroinflammatory cytokines that had the greatest response to Tat-Cocaine are shown, with this treatment resulting in increased levels of IL-1α, IL-1β, IL-6, TNF-α, MIP-1α, MCP1, and LIX cytokines. 4OI treatment reduced the release of these cytokines by at least 30% (Figure 5). In particular, the levels of IL-1α and IL-1β were reduced by >50% in both the Tat-only or Tat-Cocaine treatment groups in the presence of 4OI. This data clearly shows that 4OI treatment suppresses the pro-inflammatory cytokine release profile of mixed cortical cultures.

**Figure 5:**
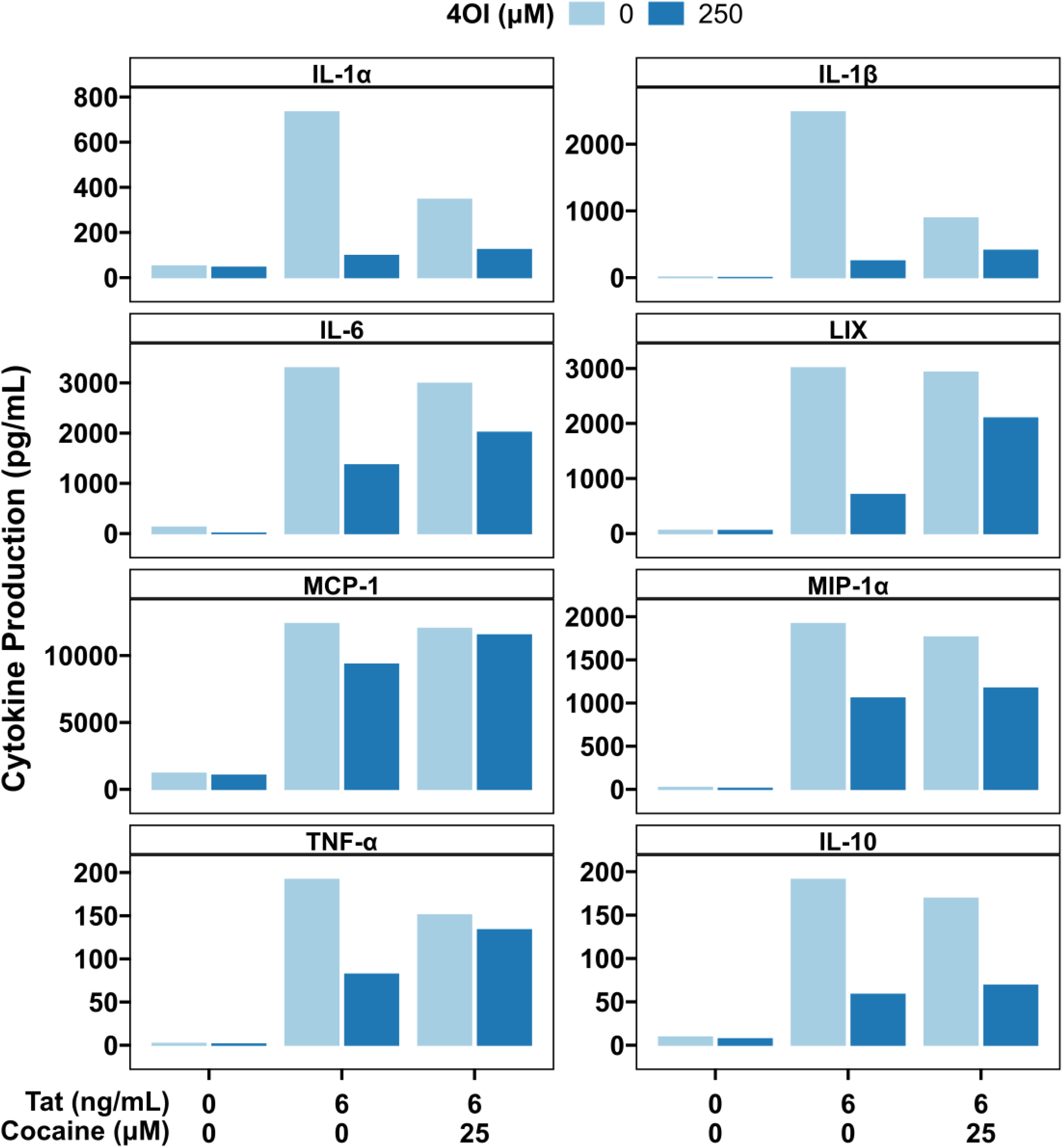
Effects of 4OI on cytokine secretion profile in primary neuronal cultures treated with Tat and/or cocaine. Bar graphs showing the chemokine/cytokine secretion levels followed by Tat (6 ng/mL) and Tat-Cocaine (25 μM), with or without 4OI (250 μM). The cytokine levels were determined by 27-plex chemokine/cytokine array from cell culture media collected from each sample. Bars represent cytokine concentrations in the collected medium (pg/ml).

Interestingly, the induction of IL-10 cytokine levels followed the exposure to Tat or Tat-Cocaine, which was notably suppressed with 4OI treatment. Although traditionally recognized as an anti-inflammatory cytokine, emerging evidence suggests that IL-10 can act as a modulator contributing to the formation and endurance of HIV reservoirs in non-primate HIV models (Harper, Ribeiro et al. 2022). Our cell culture model aligns with and supports this evidence.

## DISCUSSION

Previously, we developed and applied a literature mining and transcriptomics-based approach to uncover new compounds that can inhibit neurotoxic outcomes linked to both HIV and drugs of abuse, with the ultimate goal of identifying potential treatments for HAND with concurrent substance use disorders (Sybrandt, Shtutman et al. 2017, Aksenova, Sybrandt et al. 2019, Sybrandt, Tyagin et al. 2020). The analysis of transcriptomes involved a well-established model that simulates HAND with comorbid cocaine use disorder (CUD), utilizing primary cortical cultures treated with Tat and cocaine (Turchan, Anderson et al. 2001, Nath, Hauser et al. 2002, Aksenov, Aksenova et al. 2006, Midde, Huang et al. 2013, Bertrand, Hu et al. 2015, Aksenova, Sybrandt et al. 2019). This analysis revealed a significant upregulation in the expression of *Acod1* (aconitate decarboxylase 1) mRNA upon exposure to both Tat and Tat-Cocaine treatment.

The ACOD1 protein catalyzes the conversion of the tricarboxylic acid cycle intermediate cis-aconitate into itaconate, an immunometabolite produced during inflammatory responses. Itaconate exhibits potent anti-inflammatory activity (O’Neill and Artyomov 2019) and notably triggers the activation of the anti-inflammatory transcription factor Nrf2 via cysteine thiol alkylation of Keap1, inhibiting Keap1-mediated degradation of Nrf2 (Mills, Ryan et al. 2018). Further, itaconate establishes tolerance in macrophages exposed to LPS-induced inflammation, hindering late NLRP3 inflammasome activation, IL-1β release, and pyroptosis by dicarboxypropylation of GSDMD (Gasdermin D) (Bambouskova, Potuckova et al. 2021), the main regulator of NLRP3-dependent pyroptosis (Shi, Zhao et al. 2015), as well as suppresses IL-1α and IL-1β secretion (He, Wan et al. 2015, Tsuchiya, Hosojima et al. 2021). Consequently, itaconate orchestrates the regulation of tolerance and the overall level of inflammatory response in macrophages.

Given itaconate’s widely recognized anti-inflammatory properties and its potential as a therapeutic candidate for various conditions like sepsis, pulmonary inflammation, lupus, and multiple sclerosis (Lin, Ren et al. 2021), we conducted experiments to investigate the possibility of inducing an itaconate-mediated anti-inflammatory response in our HAND-CUD model by utilizing a cell-penetrable analog of this metabolite. Indeed, the ester-modified 4OI showed a potent anti-inflammatory activity in this model. Note that 4OI is also capable of penetrating the blood-brain barrier.

Using primary cortical cultures, we observed a significant increase in cell death when Tat and cocaine were administered together, indicating the synergistic and detrimental neurotoxic effects of this combination (Figure 1C-B, Supplemental Figure 1). However, 4OI treatment robustly rescued cellular survival. Excitingly, 4OI was shown to inhibit neuronal apoptosis triggered by Tat and cocaine treatment in a dose-dependent manner (Figure 1C-D). This suggests that 4OI has the ability to mitigate the toxic effects induced by Tat and cocaine and consequently could be a potential brain-penetrant neuroprotective agent.

Given the specific association of *Acod1* expression with inflammation-induced microglia (Sousa, Golebiewska et al. 2018), we subsequently proceeded to determine the effects of 4OI on microglia. Pathological stimuli are known to trigger changes in microglial morphology, which is closely linked to their functional states. These states can be categorized as "ramified resting" and "activated ameboid" (Karperien, Ahammer et al. 2013), with various intermediate forms in between. When exposed to detrimental environmental conditions, initially ramified and quiescent microglial cells transition into an intermediate form, characterized by reduced arborization and enlarged soma. Eventually these cells adopt an amoeboid morphology, involving process retraction and increased cell body sizes, which indicates enhanced motility and activity (Ekdahl 2012, Walker, Nilsson et al. 2013, Davis, Salinas-Navarro et al. 2017).

Tat and cocaine treatment revealed a concentration-dependent impact on microglia morphology, including an increase in microglial cell number, a shortening of the microglia processes, and an increase in microglia size, with the exception of the highest tested combination concentration, at which survival was substantially compromised (Figure 3A-C). The percentage of Iba1-positive cells within the culture decreased, and microglia transitioned into a dystrophic state, characterized by spheroidal swelling and stripped ramifications and processes (Holloway, Canty et al. 2019) (Figure 3A-C). Although it appeared by morphology that 4OI and Tat, Tat/cocaine treatments all shift microglial morphology to ameboid, the RNA-seq results suggest that 4OI treatment results in a non-inflammatory ameboid microglia, which have been reported in normal neonatal and adult brains (Orlowski, Soltys et al. 2003, Silva, Dorman et al. 2021)

In line with previous observations, 4OI treatment led to a significant activation of Nrf2-regulated antioxidant genes (Saha, Buttari et al. 2020, Simpson and Oliver 2020, Peace and O’Neill 2022) in both untreated primary cortical cultures and those exposed to Tat and cocaine (Figure 4). Furthermore, 4OI treatment suppressed the activation of pro-inflammatory pathways specific to myeloid cells, including ones like the adaptive immune response and complement binding. These pathways, typically set in motion by Tat and Tat-Cocaine, are known to be activated by HIV proteins through Toll-like receptor dependent pathways (Donninelli, Gessani et al. 2016). In agreement with RNA-seq data, the secretion of proinflammatory cytokines such as IL-1α, IL-1β, IL-6, TNF-α and MIP-1α were activated by Tat and Tat-Cocaine and suppressed by 4OI.

Interestingly, the effects of 4OI on the number of microglial cells were context dependent. While 4OI treatment increased the number of cells in control and Tat-treated cultures, 4OI conversely decreased the amount of microglial cells in Tat-Cocaine exposed cultures (Figure 3). These results could be attributed to the previously-documented cell- and condition-specific effects associated with Nrf2 activation on cellular proliferation, which can yield either pro- or anti-proliferative effects (Murakami and Motohashi 2015). The cell and condition specific contexts that contribute to pro- or anti-proliferative effects in our model system require further investigation in future studies.

Moreover, 4OI treatment countered the Tat and Tat-Cocaine dependent upregulation of Phosphodiesterases (*Pde4b* and *Pde5a*) and Purinergic receptors (*P2ry2*, *P2ry6* and *P2rx4*). Phosphodiesterases, especially those of *Pde4* and *Pde5* families, are well-known regulators of microglia activation and serve as targets for the development of small molecules to manage neuroinflammation and treat neurodegenerative diseases, alongside other neuroinflammation-related pathologies (Pearse and Hughes 2016, Sanders 2020). The microglial purinergic receptors are activated by nucleoside di- and triphosphates released by neurons and glial cells in response to both normal brain activity and as a consequences of pathological events (Calovi, Mut-Arbona et al. 2019). Specifically, receptors like P2ry2 and P2ry6, located on the surface of inflammatory microglia, play a role in the induction of phagocytosis and in the removal of stressed neurons (Puigdellivol, Milde et al. 2021, Hide, Shiraki et al. 2023). The expression of P2Y receptors is regulated in concert with the genes governing phagocytosis, such as *VASP* (Montaño-Rendón, Walpole et al. 2022), *Rac2* (Flannagan, Canton et al. 2014) and *TGM2* (Rébé, Raveneau et al. 2009).

Strikingly one of the most downregulated genes in Tat and Tat-Cocaine treated cultures by 4OI was *Acod1*. This result points to a potential autoregulatory mechanism of *Acod1* expression by its product itaconate that balances activation of itaconate-dependent anti-inflammatory signals in microglial cells (Figure 6). However, it is also possible that the effects of 4OI could differ from those of endogenous itaconate (Swain, Bambouskova et al. 2020). Thus, additional studies are needed to substantiate the presence of such an autoregulatory loop.

**Figure 6:**
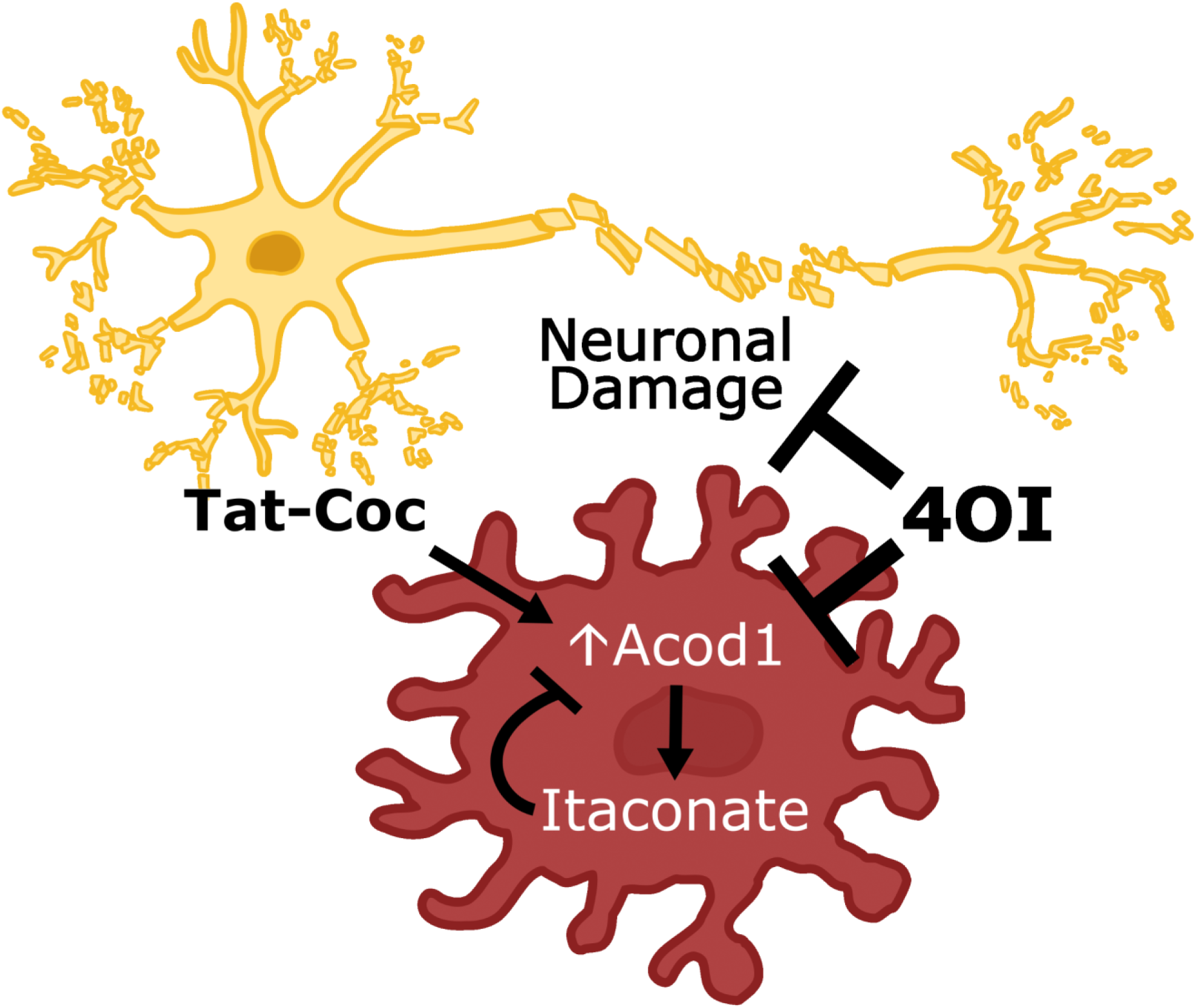
Possible model of 4OI-mediated neuroprotection. The observed neuronal damage and Acod1 overexpression resulting from Tat and cocaine treatment are significantly attenuated upon 4OI administration. This potential neuroprotection might stem from 4OI’s negative feedback inhibition of Acod1

Collectively, these findings represent the first exploration of the role of cell-penetrable itaconate esters in microglia activation within the context of HAND-CUD models. Although the precise mechanism of itaconate-dependent microglia activation and regulation requires further studies, the application of the cell- and brain-penetrable form of itaconate, 4OI, holds promise as a potential therapeutic approach for HAND. Furthermore, investigations are warranted to elucidate the underlying mechanisms and evaluate the translational potential of 4OI in treating neuroinflammatory disorders and other neurodegenerative diseases.

## List of abbreviations

ART: antiretroviral therapy
PLWH: people living with HIV
HAND: HIV-associated neurocognitive disorder
CUD: cocaine use disorder
BBB: blood brain barrier
CNS: central nervous system
IRG1: immune responsive gene 1
Acod1: aconitate decarboxylase 1
4OI: 4-octyl-itaconate
Nqo1: NAD(P)H quinone oxidoreductase 1
Gstp1: Glutathione S-transferase Pi
Gclc: Glutamate cysteine ligase catalytic subunit
VASP: Vasodilator Stimulated Phosphoprotein

## Acknowledgments

We thank COBRE Center for Targeted Therapeutics, Microscopy and Flow Cytometry Core for image analysis and Functional Genomics Core for transcriptomics analysis. The COBRE CTT cores are supported by NIH NIGMS P20GM109091. The work was supported by awards from NIH NIDA R21DA058586 (MS, RB, NF), R01DA054992 (MS, MDW).

## Figure Legends

*Supplemental Figure 1: Effects of Tat and/or cocaine on cell viability and microglia morphology in primary cortical cultures*

**(A)** Boxplots showing the percentage of dead/apoptotic cells in primary cortical cultures treated with Tat (1, 6, or 60 ng/mL) for 48 h and/or cocaine (10 or 25 μM) for an additional 24 h. The x-axis represents increasing Tat concentration, and the colors indicate different cocaine concentrations (Orange: 10 μM; Red: 25 μM).

**(B)** Percentage of Iba1-positive microglia cells in each experimental group. Each data point corresponds to an image containing 300-450 cells.

**(C)** Quantification of the length of microglia processes.

**(D)** Measurement of microglia cell body size. Each box covers 50% of the data, and the line inside indicates the median. Statistical significance was determined using the Mann-Whitney-Wilcoxon test, followed by Bonferroni post hoc test (ns, not significant; *, p<0.05; **, p<0.01; ***, P<0.001).

*Supplemental Figure 2: Effects of Tat and cocaine on microglia cell morphology in primary cortical cultures*

**(A)** Proportion of Iba1 positive microglial cells in cultures treated with high Tat (60 ng/mL) and/or 4OI (60 or 250 µM) for 48 h, followed by an additional 24 h with or without cocaine (25 µM).

**(B)** Length of microglia processes and (**C**) microglia cell body size in the same cultures. The intensity of the blue color indicates increasing 4OI concentration (60, 250 µM). The box covers 50% of the data in each condition, and the line inside indicates the median. The Mann-Whitney-Wilcoxon test was conducted to calculate the statistical significance, followed by Bonferroni post hoc test (ns, not significant; *, p<0.05; **, p<0.01; ***, P<0.001).

*Supplemental Figure 3: The effect of 4OI treatment on Acod1 expression levels upon Tat or Tat-Cocaine treatment*

**(A)** Acod1 expression level, normalized to counts per million reads, after 48h of Tat treatment, with or without 4OI, and with or without cocaine exposure for 24 h.

**(B)** qPCR analysis demonstrating the fold change of Acod1, normalized to ACTB, following 48 h Tat, with or without 4OI, and with or without an additional cocaine exposure for 24 h.

*Supplemental Figure 4: The effect of 4OI treatment on transcriptomic profiles and pathways upon Tat or Tat-Cocaine treatment*

Dot plot of gene enrichment analyses of the differentially expressed genes in cells treated with Tat with or without 4OI for 48 h, with or without an additional 24 h of cocaine exposure. Gene ontology terms are labeled with name, sorted according to adjusted p-value and gene ratio.

*Supplemental Table 1: Results of RNA-seq transcriptomics analysis, reads count and differential expression analysis*

Column descriptions are in the worksheet “Description”. Complete results are in the worksheet “All_Data”, Results for the genes expressed differentially in any contrast (filtered by FDR < 0.05) are in the worksheet “FDR < 0.05”.

