## Supplementary figures and images for "Suppression of HIV and cocaine-induced neurotoxicity and inflammation by cell penetrable itaconate esters"

### Supplemental Figures

#### Supplemental Figure 1

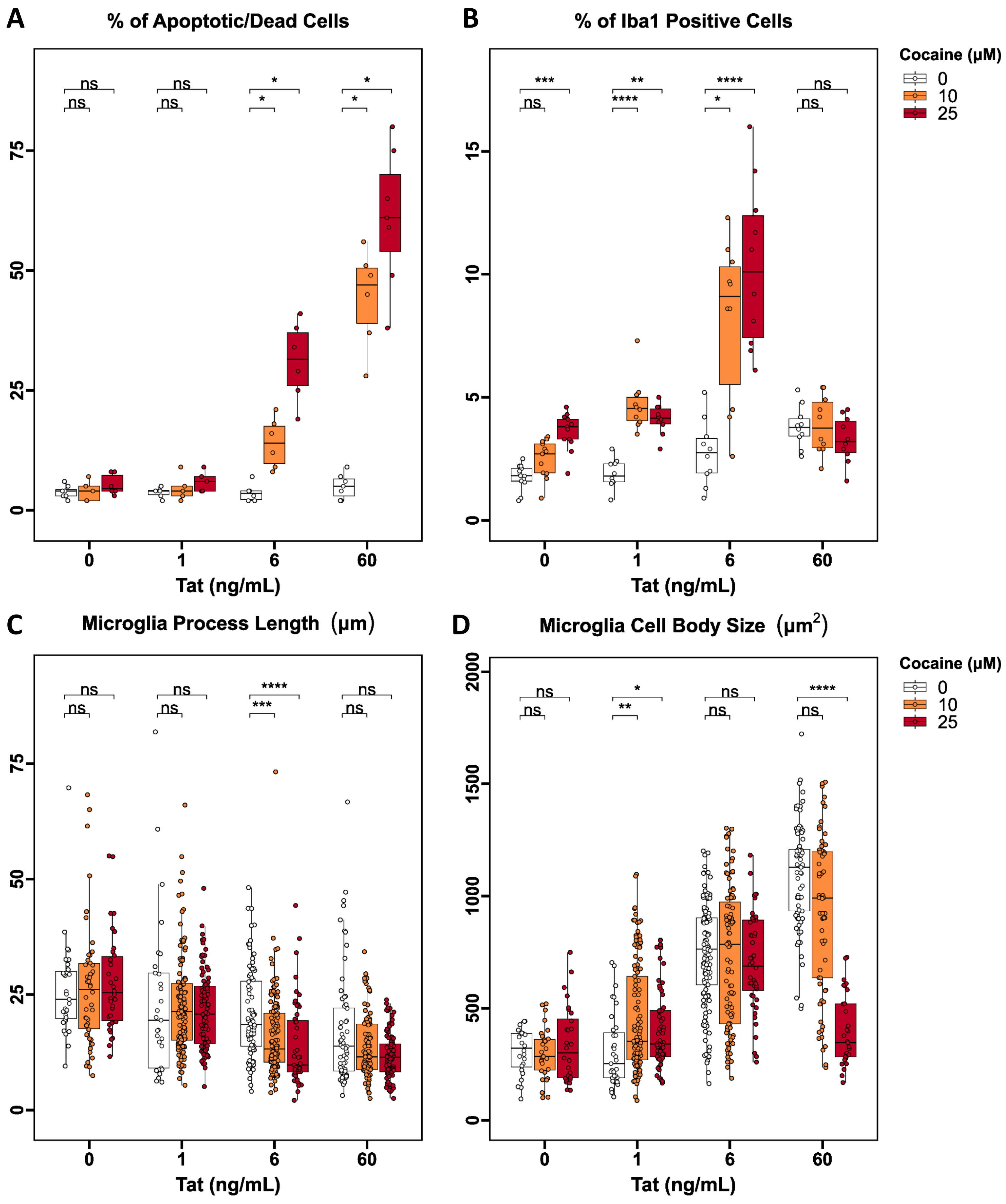

#### Supplemental Figure 2

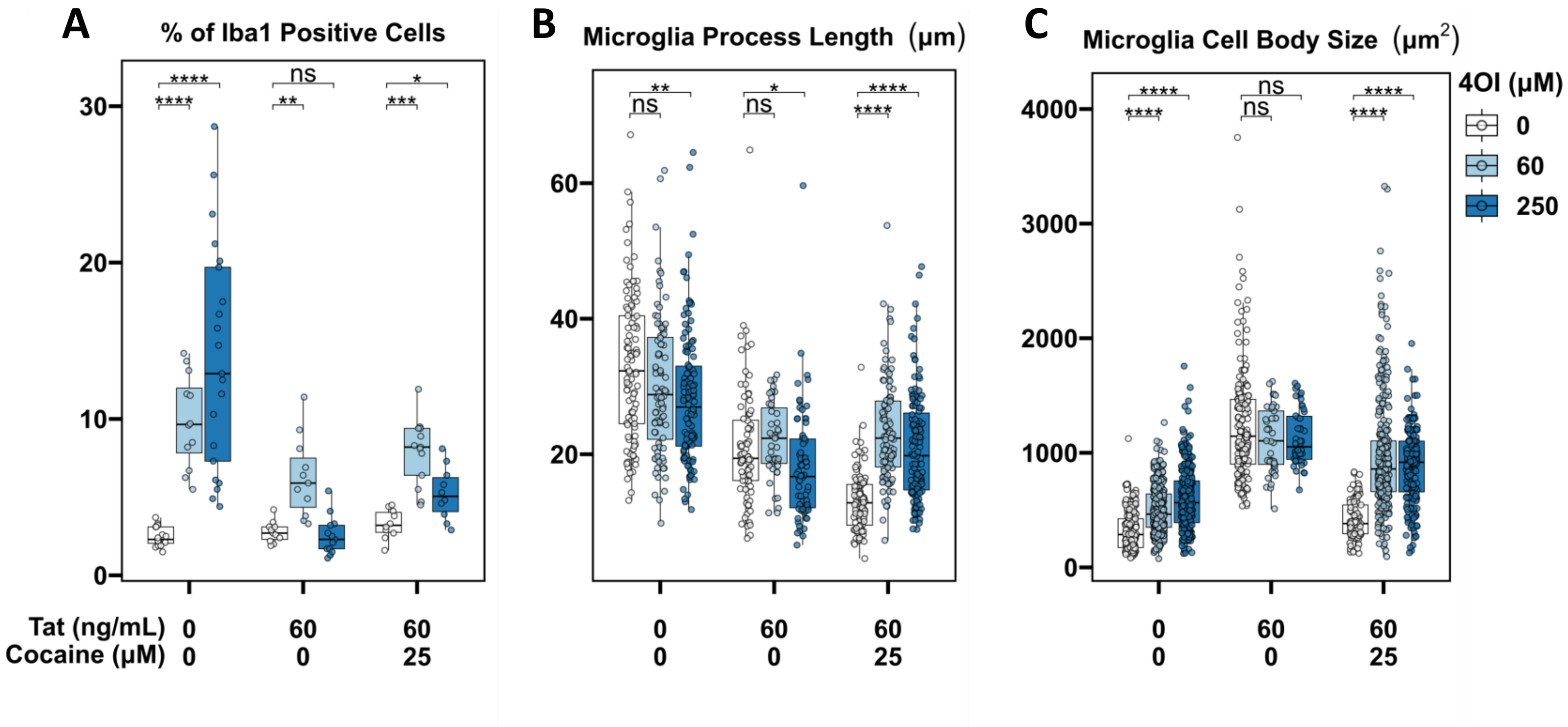

#### Supplemental Figure 3

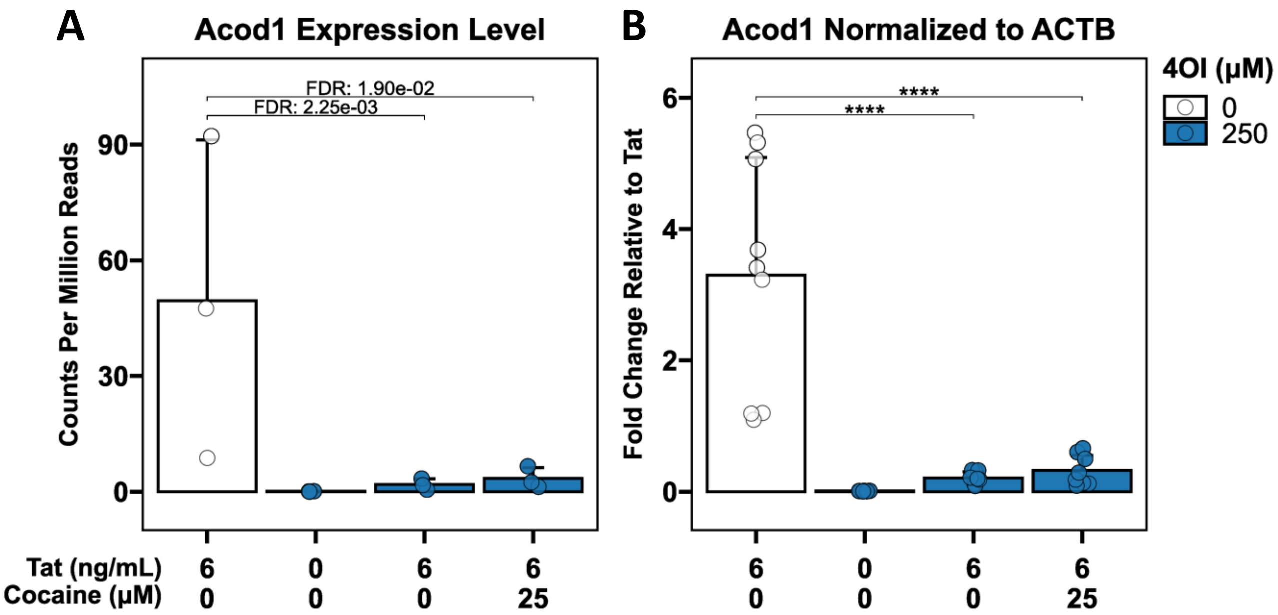

#### Supplemental Figure 4

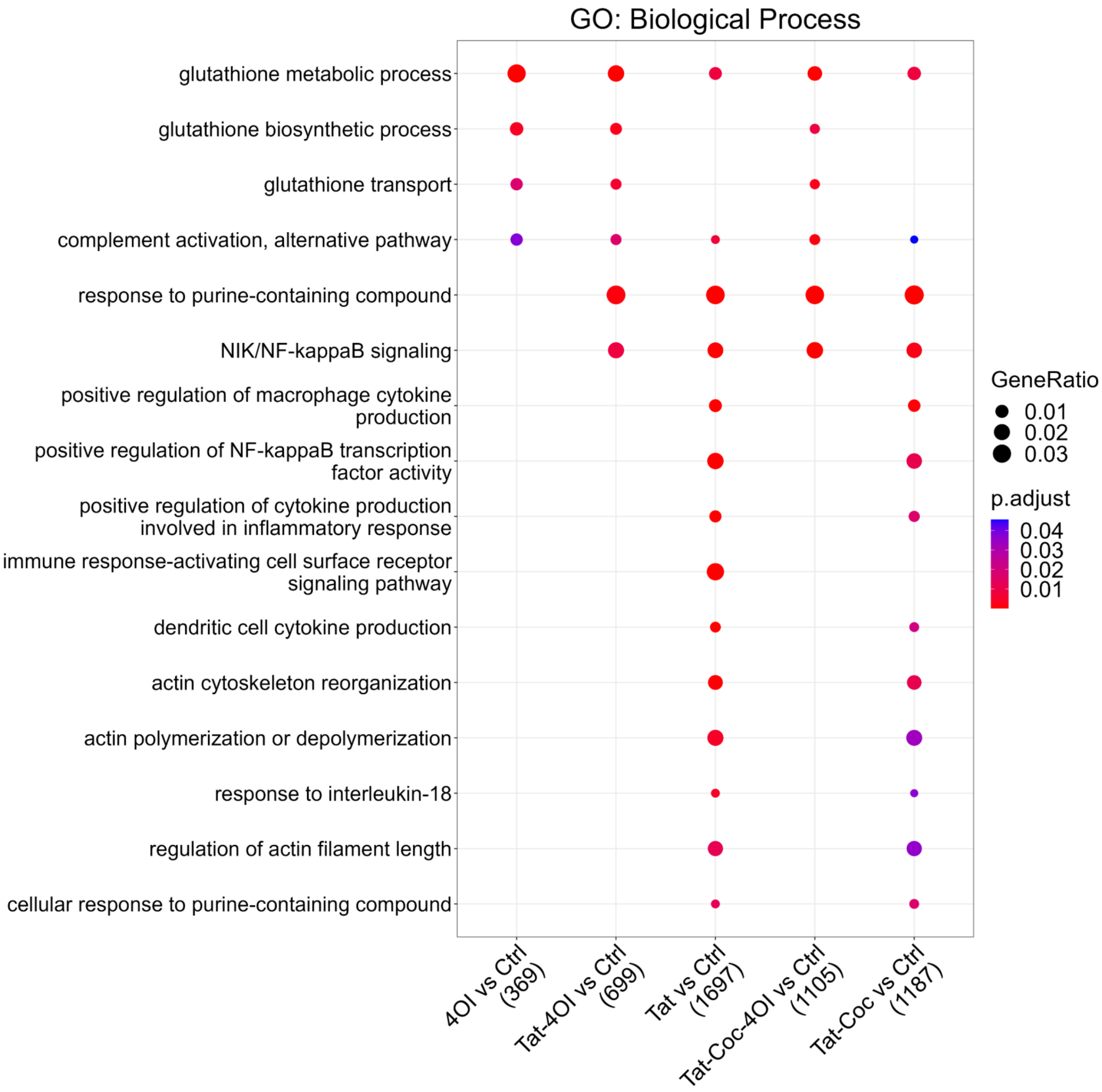
